# Matching maternal and paternal experiences underpin molecular thermal acclimation

**DOI:** 10.1101/2023.09.29.560260

**Authors:** L.C. Bonzi, J.M. Donelson, R.K. Spinks, P.L. Munday, T. Ravasi, C. Schunter

## Abstract

The environment experienced by one generation has the potential to affect the subsequent one through non-genetic inheritance of parental effects. Since both mothers and fathers can influence their offspring, questions arise regarding how the maternal, paternal and offspring experiences integrate into the resulting phenotype. We aimed to disentangle the maternal and paternal contributions to transgenerational thermal acclimation in a reef fish, *Acanthochromis polyacanthus*, by exposing two generations to elevated temperature (+1.5°C) in a full factorial design and analyzing the F2 hepatic gene expression. Paternal and maternal effects showed common but also parent-specific components, with the father having the largest influence in shaping the offspring transcriptomic profile. Fathers contributed to transgenerational response to warming through transfer of epigenetically controlled stress-response mechanisms while mothers influenced increased lipid metabolism regulation. However, the key to acclimation potential was matching thermal experiences of the parents. When both parents were exposed to the same condition, offspring showed increased structural RNA production and transcriptional regulation, whereas environmental mismatch in parents resulted in maladaptive parental condition-transfer, revealed by translation suppression and endoplasmic reticulum stress. Interestingly, the offspring’s own environmental experience had the smallest influence on their hepatic transcription profiles. Taken together, our results show the complex nature of the interplay between paternal, maternal and offspring cue integration, and reveal that acclimation potential to ocean warming depends not only on maternal and paternal contributions, but importantly on congruent parental thermal experiences.

## Introduction

Parents can affect the phenotypes of their offspring through a range of environmentally induced non-genetic inheritance mechanisms (Bonduriansky & Day, 2009). As a consequence, the phenotype of the current generation can be influenced by the environmental experiences of the previous ones as a result of environmentally induced parental effects. The term “parental effects” has long been used as a synonym of maternal effects, as mothers were thought to be the most likely parent responsible for the transmission of non-genetic material to the offspring (Crean & Bonduriansky, 2014; Mousseau & Fox, 1998), mainly through egg provisioning of nutrients, hormones, and mitochondria. However, epigenetic mechanisms of inheritance, such as DNA methylation, histone modifications, and small non coding RNAs, have been found to also be males’ attributes (Immler, 2018), and fathers have increasingly been recognized as important in the transmission of environmentally induced parental effects (Rutkowska, Lagisz, Bonduriansky, & Nakagawa, 2020).

The existence of both maternal and paternal effects raises the question on how they are combined in their offspring’s phenotype (Bell & Hellmann, 2019). Perceived environmental cues could simply have additive effects and integrate to modulate the strength of the offspring phenotypic response. In sheepshead minnow *Cyprinodon variegatus*, for example, offspring growth is highest when paternal, maternal and offspring temperature match (Chang, Lee, & Munch, 2021). Conversely, maternal and paternal effects could interact with each other in more complex ways (Lehto & Tinghitella, 2020; Moschilla, Tomkins, & Simmons, 2022). In the freshwater snail *Physa acuta*, for example, offspring are bigger only when the mother is exposed to predatory risk, but not when both parents are (Tariel, Plénet, & Luquet, 2020), while in the beetle *Tribolium castaneum* offspring are more starvation tolerant if one but not both parents experienced cold stress (Gilad & Scharf, 2019). Males and females can indeed influence their progeny phenotypes in independent ways, either because of differences in transmission mechanisms between sexes, or because of asymmetrical investments in reproduction and parental care. As a consequence, the same environmental cue can result in parents-specific effects, like in predator-exposed mother and father sticklebacks (*Gasterosteus aculeatus*) inducing different behaviors and brain transcriptional responses in their offspring (Hellmann, Bukhari, Deno, & Bell, 2020), or paternal effects in immunity transfer and gene expression being dominant compared to the maternal ones in sex-role reversed pipefish (*Syngnathus typhle*; Beemelmanns & Roth, 2016). Finally, how parental cues are integrated in the offspring phenotype can also depend on the offspring’s own environmental experience, as well as on its sex, as daughters and sons may be differently affected or affected more prevalently by one parent only (Metzger & Schulte, 2016; Schwanz, Crawford-Ash, & Gale, 2020; Seebacher, Bamford, & Le Roy, 2023). To shed light onto the complexity of non-genetic inheritance, elaborate experimental designs are needed, in order to disentangle the different parental effects of mothers and fathers and further our understanding of their potential role in acclimation to environmental change.

If adaptive, non-genetic inheritance through environmentally induced parental effects might enable acclimation to persistent environmental changes, such as global warming (Donelan et al., 2020; Donelson, Salinas, Munday, & Shama, 2018). In a coral reef fish, the spiny chromis *Acanthochromis polyacanthus* (Bleeker 1855), for instance, studies have shown that parental developmental exposure to elevated temperature can provide transgenerational acclimation to ocean warming and restoration of aerobic scope in offspring through molecular changes in lipid and carbohydrate metabolism, as well as immune system and transcriptional regulation (Bernal, Ravasi, Rodgers, Munday, & Donelson, 2021; Bernal, Schmidt, Donelson, Munday, & Ravasi, 2022; Bonzi et al., 2023; Donelson, Munday, McCormick, & Pitcher, 2012; Ryu, Veilleux, Donelson, Munday, & Ravasi, 2018; Veilleux et al., 2015). The relative importance of fathers and mothers in such acclimation process, however, is unknown, but has implications on adaptive potential. Parental pairs, for instance, might be formed by individuals differently exposed to marine heatwaves (Frölicher, Fischer, & Gruber, 2018), resulting in mothers and fathers having divergent past thermal experience and thus potentially imparting different parental effects. Understanding how these multiple signals from maternal, paternal and personal experiences are integrated into the resulting phenotype will help make predictions about population acclimation potential in a changing world.

In this study, we investigate the molecular processes underlying paternal and maternal effects of ocean warming in *A. polyacanthus* and how they interact with each other, as well as the thermal experience of the offspring. We used a full factorial, split clutch design where we developmentally exposed *A. polyacanthus* females, males or both sexes, as well as their offspring to either control or elevated (+1.5°C) temperature (Fig. 1), and then analyzed the offspring hepatic gene expression profiles. Using fish from this same experiment, Spinks et al. (2021) found parent-specific effects in life history traits and physiological performance, with mothers producing larger eggs and better quality offspring when exposed to warming, while fathers sired fewer and poorer quality offspring. There was also a non-additive interaction between maternal and paternal effects on offspring swimming performance, with fish from mismatched parents swimming faster than offspring of parents that developed at the same temperature (Spinks, 2021). Here we aim to: 1) tease apart the molecular processes regulating the individual maternal and paternal effects associated with acclimation to elevated temperature in juvenile fish, 2) investigate the interaction between the different parental experiences and thermal acclimation to elevated temperature in juvenile fish, and 3) unravel how the paternal, maternal and offspring experiences of warming integrate in the resulting molecular phenotype. Overall, with this study we aim to establish key molecular mechanisms involved in non-genetic inheritance and the relative importance of each parent in transgenerational acclimation to ocean warming.

**Figure 1.**
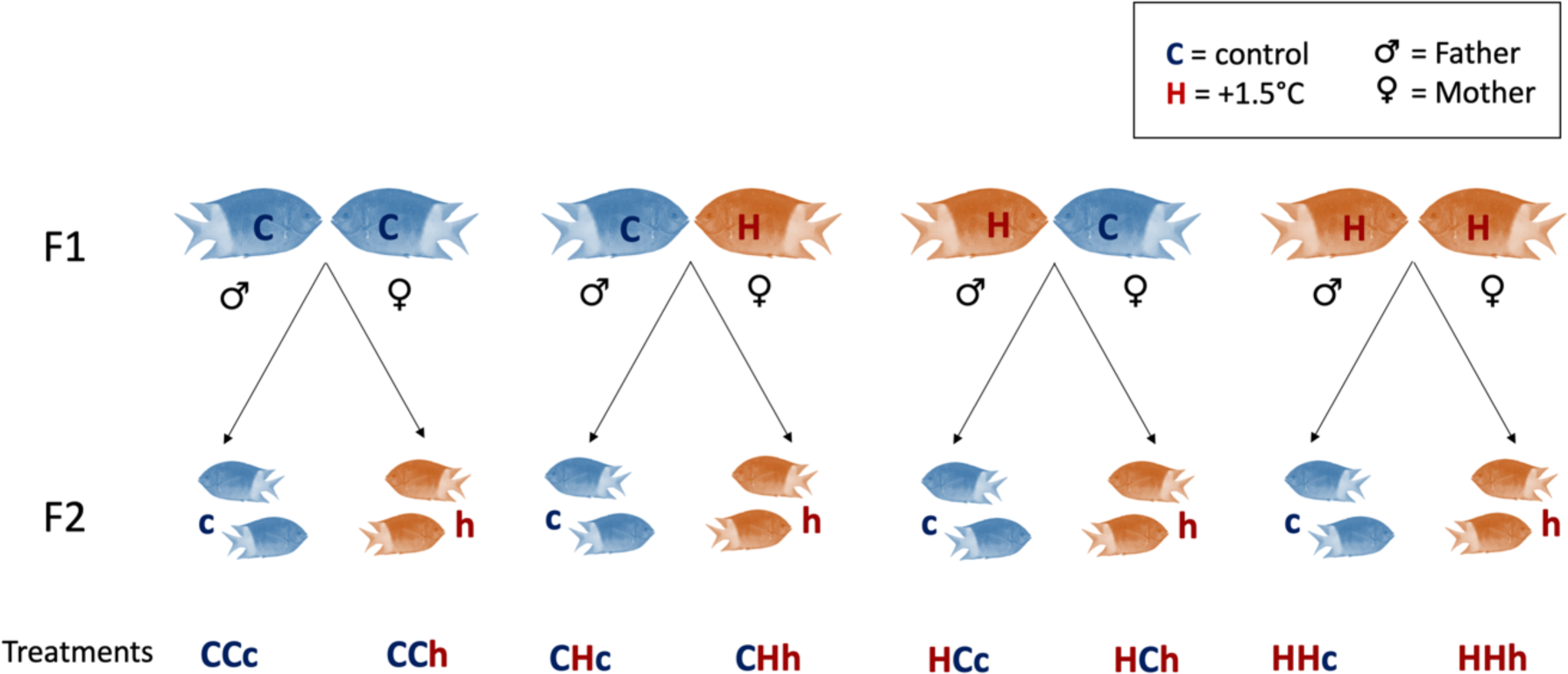
Experimental design. F1 males and females developed either at present-day control temperature (in summer 28.5°C with ±0.6°C diurnal variation), or at +1.5°C elevated temperature (in summer 30.0°C with ±0.6°C diurnal variation). At 1.5 years of age, F1 were paired in reciprocal sex crosses of the two thermal treatments. F2 were split after hatching into present-day control or elevated temperature, where they developed for 80 days. Treatment code: first letter = developmental thermal treatment of the F1 father; second letter = developmental thermal treatment of the F1 mother; third letter = developmental thermal treatment of F2 offspring.

## Materials and methods

### Experimental design

To investigate the parental contributions to thermal acclimation, we analyzed the liver gene expression of 80-day old F2 *A. polyacanthus* offspring produced in the experimental set-up described in Spinks et al. (2021; 2022). Briefly, wild adult spiny chromis damselfish (F0 generation) were collected from the Palm Islands region (18°37′ S, 146°30′ E) and nearby Bramble Reef (18°22′ S, 146°40′ E) of the central Great Barrier Reef (GBR), Australia, paired and housed with seasonally cycling water temperature in the Marine and Aquaculture Research Facility of the James Cook University in Townsville, Australia.

In the Austral summer of early 2016 newly hatched F1 offspring from six breeding pairs were split into two thermal treatments, present-day control and elevated temperature. Different from most past experiments on ocean warming, we included both seasonal (minimum 23.2°C in winter, maximum 28.5°C in summer) and diurnal (3:00 am −0.6°C, 3:00 pm hours +0.6°C) cycles that are representative of the Palm Islands region of the GBR in the two temperature treatments, to better mimic natural temperature variation and more accurately predict *A. polyacanthus* responses to ocean warming. The elevated thermal treatment followed the same cycles but with an increase of 1.5°C, chosen to match the projections for ocean warming by the end of the century in a low CO_2_ emission scenario (Fox-Kemper et al., 2021), as well as the already occurring heatwaves on the GBR (Frölicher et al., 2018). F1 fish developed and were reared in each of the two thermal treatments until they reached maturity at approximately 1.5 years of age. At this time, all fish were moved to present-day control temperature and F1 breeding pairs were formed between non-sibling fish, so that reciprocal sex crosses of the developmental temperatures were created. Reproduction for all pairs occurred at control temperature to ensure that observed phenotypic effects of developmental warming were not confounded by the exposure to elevated temperature during embryogenesis, since eggs are left with the parents until hatching to allow parental care. This resulted in F1 breeding pairs consisting of four pair combinations of males and females: 1) both sexes developed at present-day control temperature (CC), 2) males developed at elevated temperature and females developed at control temperature (HC), 3) males developed at control temperature and females developed at elevated temperature (CH) or 4) both sexes developed at elevated temperature (HH; Fig. 1).

In the Austral summer of 2017-2018 F1 breeding pairs produced the F2 generation. The first clutch of each parental pair (three to five clutches per parental treatment in total) was split into two replicate tanks of 20 fish each per thermal treatment, either present-day control or elevated temperature, which followed the above-mentioned seasonal and diurnal cycles. This resulted in a full factorial, split clutch design of a total of eight different final treatments (Fig. 1). F2 fish were reared in the described thermal treatments until 80 days post-hatching (dph), when they were sexed by urogenital papilla external examination, and two males and two females per clutch (twelve to twenty individual fish per treatment) were euthanized by cervical dislocation, measured for length, weighed for mass and dissected. Livers were immediately snap-frozen in liquid nitrogen and stored at −80°C for subsequent RNA extraction. All samples were collected between 9 am and 12 pm. All procedures were performed in accordance with relevant guidelines and were conducted under James Cook University’s animal ethics approval A1990, A2210 and A2315.

### RNA sequencing and gene expression analysis

Samples were processed as already in Bonzi et al. (2023). Briefly, total RNA was isolated from homogenized whole livers using a *mir*Vana miRNA Isolation kit. Liver was chosen because of its major role in metabolism, and to allow comparison more easily with previous works (Bernal et al., 2018; Bernal et al., 2021; Veilleux et al., 2015). Isolated RNA was checked for quality and quantity, and mRNA-focused libraries were prepared using the Illumina TruSeq stranded mRNA Library Preparation Kit. Libraries were paired-end sequenced with Illumina HiSeq 4000 (150 bp) at the King Abdullah University of Science and Technology Bioscience Core Lab. FastQC (Andrews, 2010) was used to quality check the raw reads before and after Trimmomatic v0.39 (Bolger, Lohse, & Usadel, 2014) quality trimming and adapter removal, using a sliding window of 4:15 and retaining paired-end reads with minimum length of 40 bp. Trimmed reads were then mapped against the *Acanthochromis polyacanthus* genome (ENSEMBL ASM210954v1) using HiSAT2 v2.2.1 (Kim, Paggi, Park, Bennett, & Salzberg, 2019), and featureCounts (Liao, Smyth, & Shi, 2014) from the Subread v2.0.2 package was used to calculate gene counts, in read pair counting mode, allowing for multi-mapping fractional computation.

The resulting count matrix was then imported in R v.3.6.3 (R Core Team, 2020), where DESeq2 package v1.26.0 (Love, Huber, & Anders, 2014) was used to detect differentially expressed genes across treatments. Raw counts were variance-stabilizing transformed (VST) using the *vst* function, and principal component analyses (PCAs) and heatmaps of the sample-to-sample distances were run to check for outliers and batch effects. Based on the resulting plots, four outlier samples were excluded from further analyses, leaving a total of 124 samples (11–20 samples/treatment; Suppl. Table S1). In order to choose the best design formula for differential gene expression analyses, likelihood ratio tests (LRTs) were used to determine which variables (e.g., family, sex, size) and interactions between treatments were significant. The resulting design formula included the variable “family” to acknowledge the parental lineage, and the two-way interaction between maternal and paternal developmental thermal environments, besides the paternal, maternal and offspring developmental treatments as main effects: ∼Family + Paternal developmental temperature * Maternal developmental temperature + Offspring developmental temperature. Differentially expressed genes (DEGs) due to the parental interaction and to the main effects (padj < 0.05) were then statistically determined. Pairwise comparisons were run between the treatments using Wald tests, followed by log2 Fold Change (log2FC) shrinkage by apeglm method (Zhu, Ibrahim, & Love, 2019). The threshold cut-offs chosen to identify significant DEGs in the pairwise comparisons were False Discovery Rate (FDR) adjusted *p*-value < 0.05 (Benjamini & Hochberg, 1995), |log2FC| ≥ 0.3 to reduce false positives, and a mean expression of > 10 reads (baseMean). Finally, LRT identified DEGs due to parental treatment interaction were clustered using the *degPatterns* function from the DEGreport R package v1.22.0 (Pantano, 2021) to identify similar expression profiles. The function was run on the VST processed count matrix of such genes with default settings, except for cluster outlier removal (reduce = TRUE).

Functional enrichment analyses of differentially expressed genes and expression clusters were performed in OmicsBox v2.0.36 (OmicsBox, 2019) with the Fisher’s Exact Test (FDR < 0.05).

## Results

### Overall influence of parental and offspring thermal environments

Paternal and maternal developmental exposures to elevated temperature caused the differential regulation of a common component of genes in their offspring’s liver, as well as parent-specific sets of genes, which were overall mostly distinct from the offspring’s own thermal exposure molecular signature (Fig. 2). The offspring’s own exposure to warming resulted in 1,579 differentially expressed genes (DEGs), mainly involved in oxidoreductase activity, metabolism, DNA replication, and vacuolar transport (Fig. 3; Suppl. Table S2). Conversely, paternal exposure to elevated temperature caused the differential regulation of 6,684 genes in the offspring, while maternal exposure to warming resulted in 5,877 DEGs (Fig. 2; Suppl. Tables S3 & S4, respectively). Roughly half (3,914) of the genes influenced by parental exposure to elevated temperature were common to the warming exposure of both parents (Fig. 2) and are involved in metabolism, translation, gene expression, ncRNA processing, T cell activation, semaphorin-plexin signaling pathway and telomere maintenance (Fig. 3; Suppl. Table S5). We did not find any offspring sex-specific parental effects, or differential gene expression between male and female offspring. Testing the interaction between the two generations’ thermal treatments resulted in no DEGs, indicating that the gene expression patterns in the offspring were not influenced by the match or mismatch of their thermal experience with the parental environment. A strong interaction was instead found between the paternal and the maternal thermal experiences.

**Figure 2.**
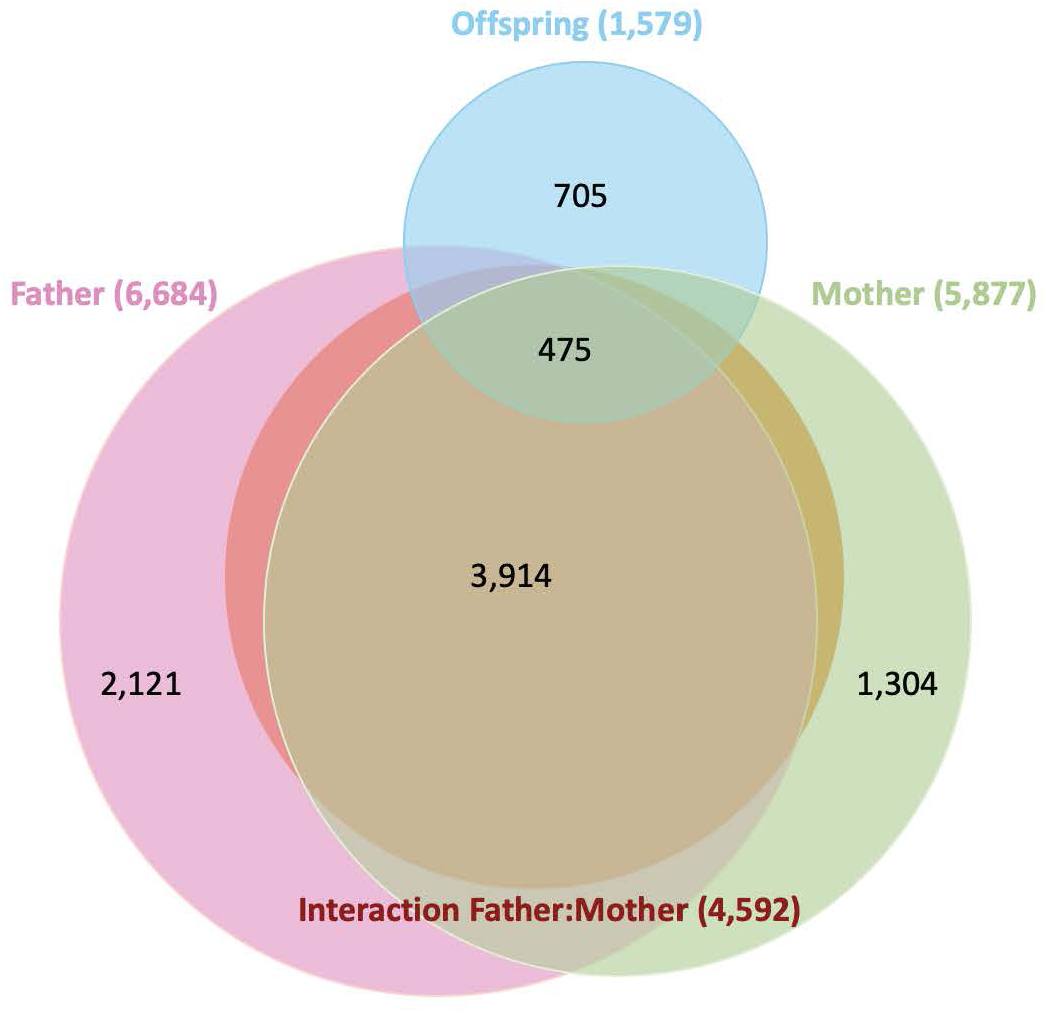
Venn diagram of differentially expressed genes attributable to the paternal, maternal, offspring developmental thermal experiences, and to the interaction between the paternal and the maternal ones, identified by likelihood ratio tests.

**Figure 3.**
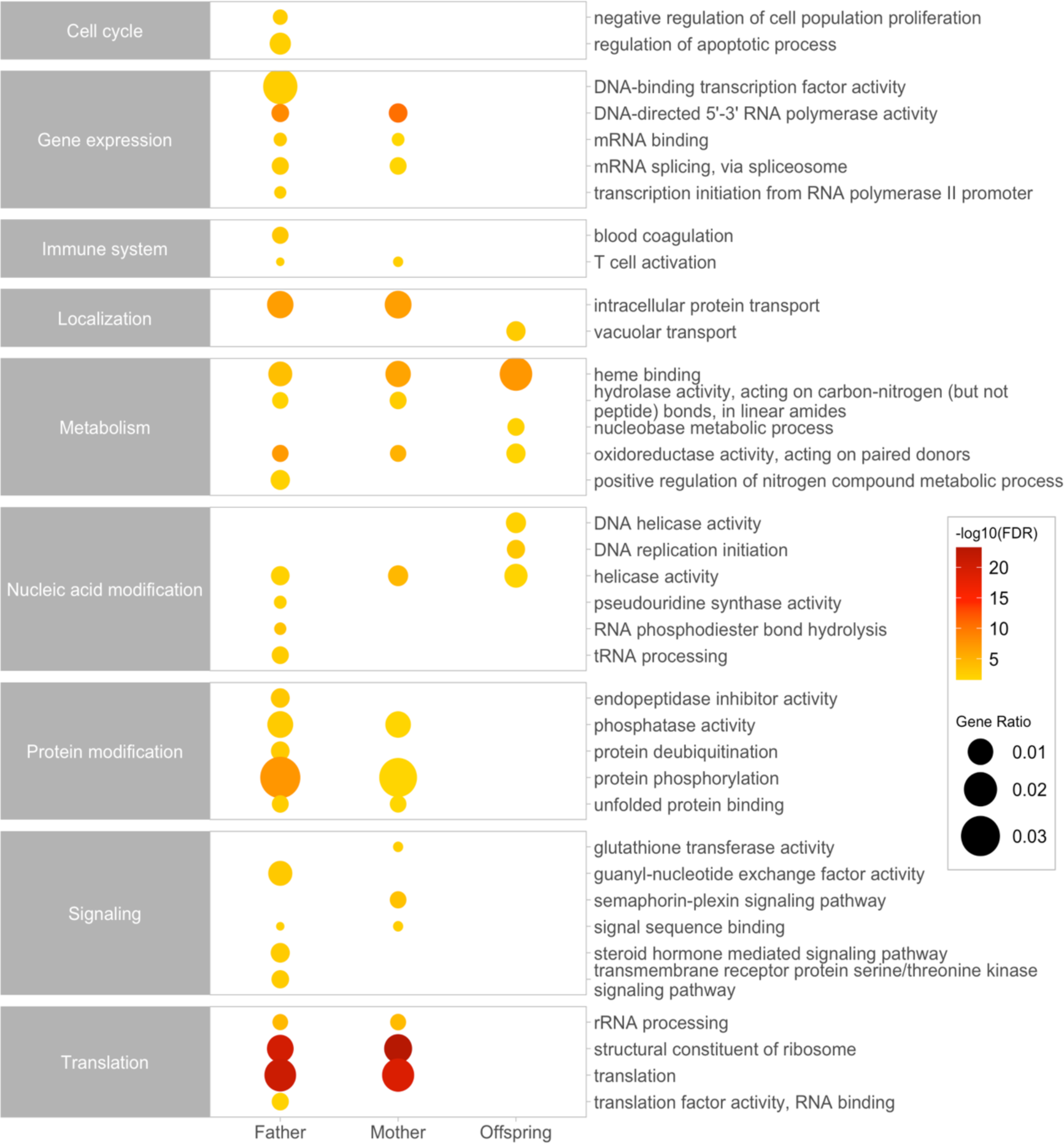
Functional enrichment analysis of differentially expressed genes attributable to the Father, Mother and Offspring developmental thermal experiences. The color of the circles represents the enrichment significance with darker coloration increasing in significance, and size of circles is proportional to the number of genes underlying the enriched function.

### Paternal thermal molecular signatures

The exposure of fathers to warming resulted in 2,121 father-specific DEGs in the offspring (Fig. 2; Suppl. Table S3). Processes involved in protein modification, RNA processing, negative regulation of cell population proliferation, apoptosis and small GTPase mediated signal transduction were overrepresented in this set of genes, as well as functions related to transcription and post-transcriptional regulation of gene expression (Fig. 3; Suppl. Table S6). The pairwise comparison between offspring from fathers exposed to warming and control mothers (HC) and offspring where both parents were exposed to control conditions (CC) resulted in 2,902 DEGs, regardless of the offspring’s own developmental temperature (Fig. 4; Suppl. Table S7). The upregulated genes in HC offspring are involved in small GTPase mediated signal transduction, transcription, regulation of apoptotic process, DNA repair, response to stress and deacetylation of proteins (e.g., histones). The processes controlled by the downregulated genes are related to translation, semaphorin-plexin signaling and chromatin remodeling via SWI/SNF complex (Suppl. Table S8). Among these DEGs, 128 were shared with the differences that were found between offspring of parents that were both exposed to warming (HH) compared to CC offspring, but not with offspring from control fathers and mothers exposed to warming (CH) compared to CC offspring (Suppl. Fig. S1A; Suppl. Table S7). This consistency in patterns of gene expression between HC vs CC and HH vs CC provides evidence for a paternal effect independent of the maternal experience. Inflammation and innate immune system were downregulated, as well as adipogenesis (AGT, BMP1, ID3, LIFR, NCOR2, STAT5B) and fatty acid metabolism, while other shared DEGs were involved in apoptosis, transcription regulation, chromatin organization and epigenetic regulation of gene expression, through histone acetylation (KANSL3, NAA40) and miRNA mediated gene silencing (TNRC6B, TNRC6C). When HC offspring developed at elevated *versus* control temperature, they differentially expressed 140 genes, with 126 DEGs exclusive to this pairwise comparison (Fig. 5; Suppl. Table S9). These genes are mainly involved in the downregulation in offspring exposed to elevated temperature of actin cytoskeleton organization and immune response. RNA processing and autophagy related genes were instead upregulated, while other genes are involved in transcription regulation, metabolism, proteolysis, and signal transduction.

**Figure 4.**
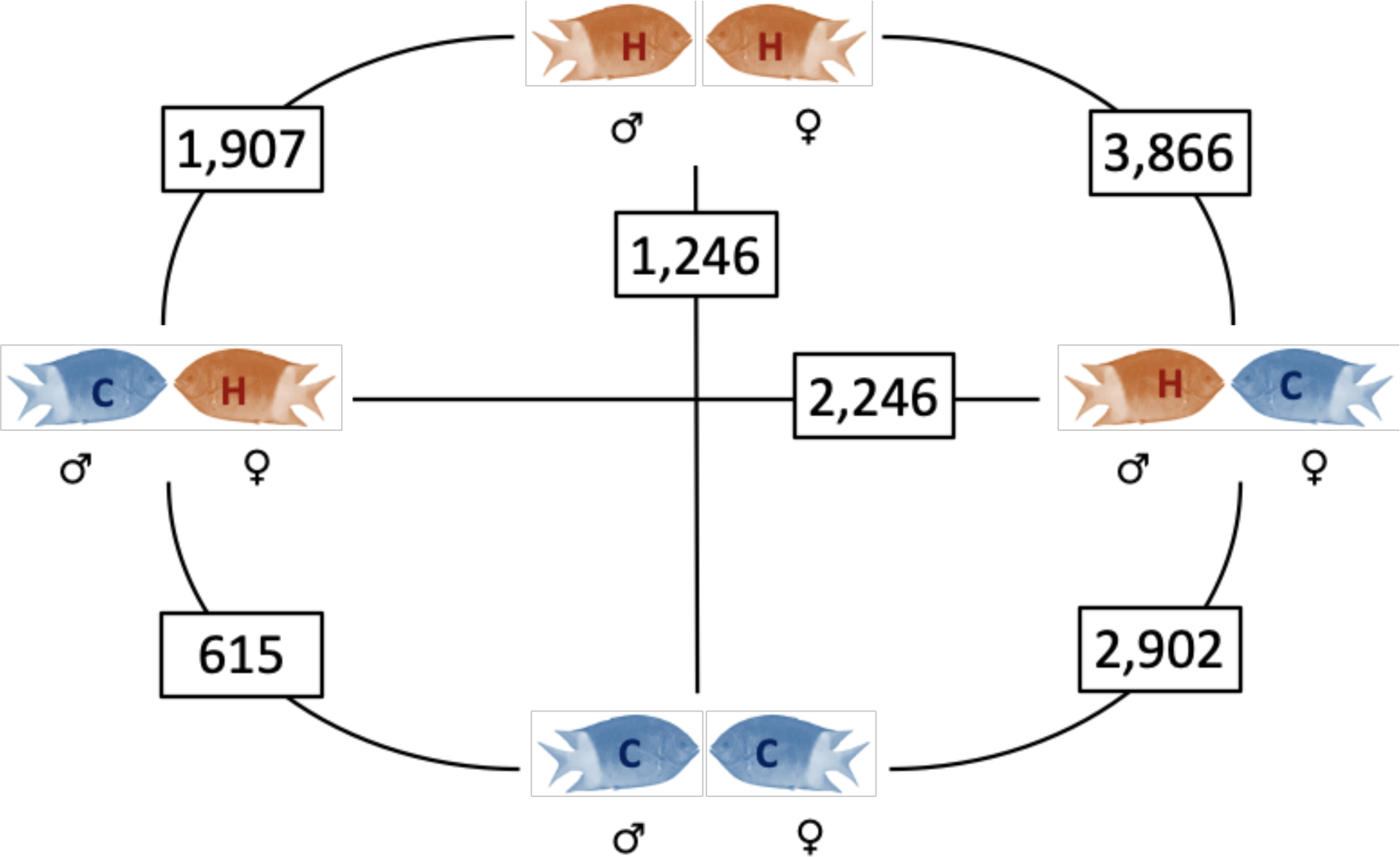
Numbers of genes differentially expressed in offspring from different parental treatments, regardless of their own developmental thermal environment. “C” stands for control temperature, “H” stands for +1.5°C.

**Figure 5.**
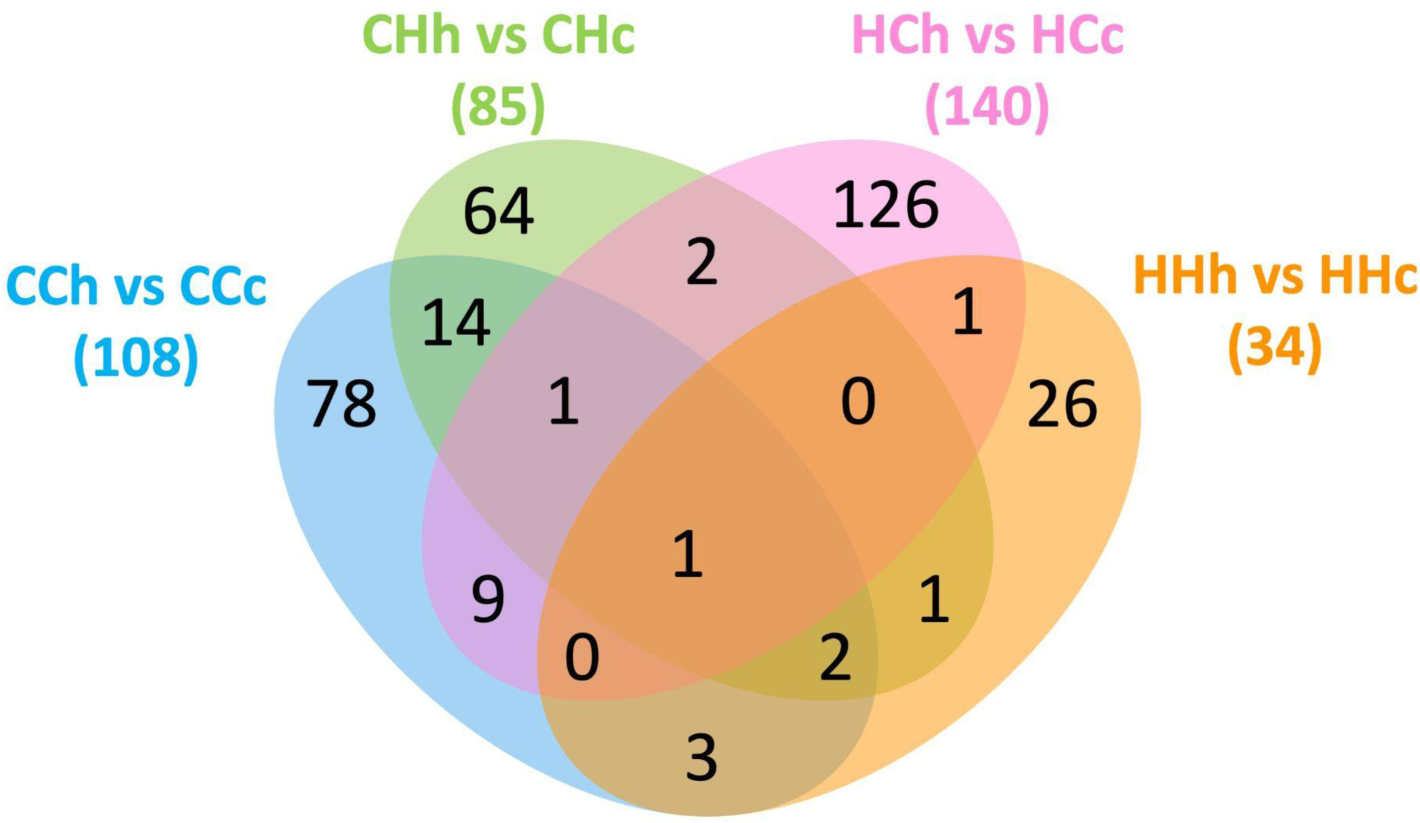
Venn diagram of differentially expressed genes in pairwise comparisons between offspring from the same parental treatments (first capital letter represents the paternal thermal environment, second capital letter represents the maternal thermal environment) developed at elevated (h) *versus* control temperatures (c; lowercase letter). “C/c” stands for control temperature, “H/h” for +1.5°C.

### Maternal thermal molecular signatures

The exposure of the mother alone to elevated temperature resulted in protein and purine ribonucleotide binding, as well as catalytic activity acting on nucleic acid and dipeptidase activity in the overall 1,304 uniquely differentially expressed offspring genes (Fig. 2; Suppl. Table S4 & S10). Offspring born from control fathers and elevated temperature mothers (CH) differentially expressed 615 DEGs compared to CC offspring, regardless of their own developmental temperature (Fig. 4; Suppl. Table S11). “Lipid transport” GO term was enriched for the upregulated genes in CH offspring, mainly due to several apolipoproteins (APOA4, APOB, APOC2, APOE), while the downregulated genes were related to processes such as glutathione transferase activity (GSTA3, GSTM3, GSTO1), structural constituent of ribosome, and telomere maintenance (Suppl. Table S12). Among these DEGs, 34 genes overlap between CH vs CC and HH vs CC comparisons, representing the maternal signature independent of the paternal or offspring thermal experiences (Suppl. Fig. S1A; Suppl. Table S11). Immune and inflammatory responses, lipid and retinol metabolism, as well as hepatocyte apoptotic process were the interested functions. CH offspring differentially expressed 64 unique DEGs compared to control (Fig. 5; Suppl. Table S13), mainly involved in metabolism of lipids and lipoproteins (ABHD3, APOA4, APOE, COQ3, CPTP, HAO2, NUS1, OSBPL10) and energy metabolism, all upregulated in offspring raised at elevated temperature.

Finally, the direct comparison between offspring from father alone (HC) *versus* mother alone (CH) exposed to warming returned 2,246 differentially expressed genes (Fig. 4; Suppl. Table S14). Compared to CH offspring, offspring from HC parents, regardless of their own thermal treatment, upregulated genes involved in small GTPase mediated signaling, oxidoreductase activity and response to stress, while downregulating genes with functions related to protein metabolism, structural constituent of ribosome, semaphorin-plexin signaling, and positive regulation of DNA-binding transcription factor activity (Suppl. Table S15).

### Interaction effects between maternal and paternal exposures to warming

Maternal and paternal exposures to warming interacted to shape the expression of 4,592 genes (Fig. 2; Suppl. Table S16). Among them, two clusters of genes are differentially regulated depending on the match or mismatch between the paternal and the maternal thermal environments (Fig. 6). Specifically, 1,861 DEGs belong to a cluster expressed at higher levels in offspring from mismatched parents (Cluster A; Fig. 6; Suppl. Table S17). Many of these genes show functions related to protein modification and transport, including members of the calnexin/calreticulin cycle and ERAD pathway involved in protein folding in the endoplasmic reticulum (ER) and involved in the response to ER stress. Other functions that characterize this group of genes are cell redox homeostasis, and lipid biosynthesis and metabolism (Suppl. Table S17). Conversely, among the 1,533 genes that followed the opposite pattern and were expressed at higher levels in offspring from matched parents (Cluster B; Fig. 6; Suppl. Table S18), we found several ribosome component genes related to translation and protein synthesis, as well as genes encoding for histones and proteins involved in gene expression, DNA-directed 5’-3’ RNA polymerase activity, transcription by RNA polymerase III and cytochrome-c oxidase activity.

**Figure 6.**
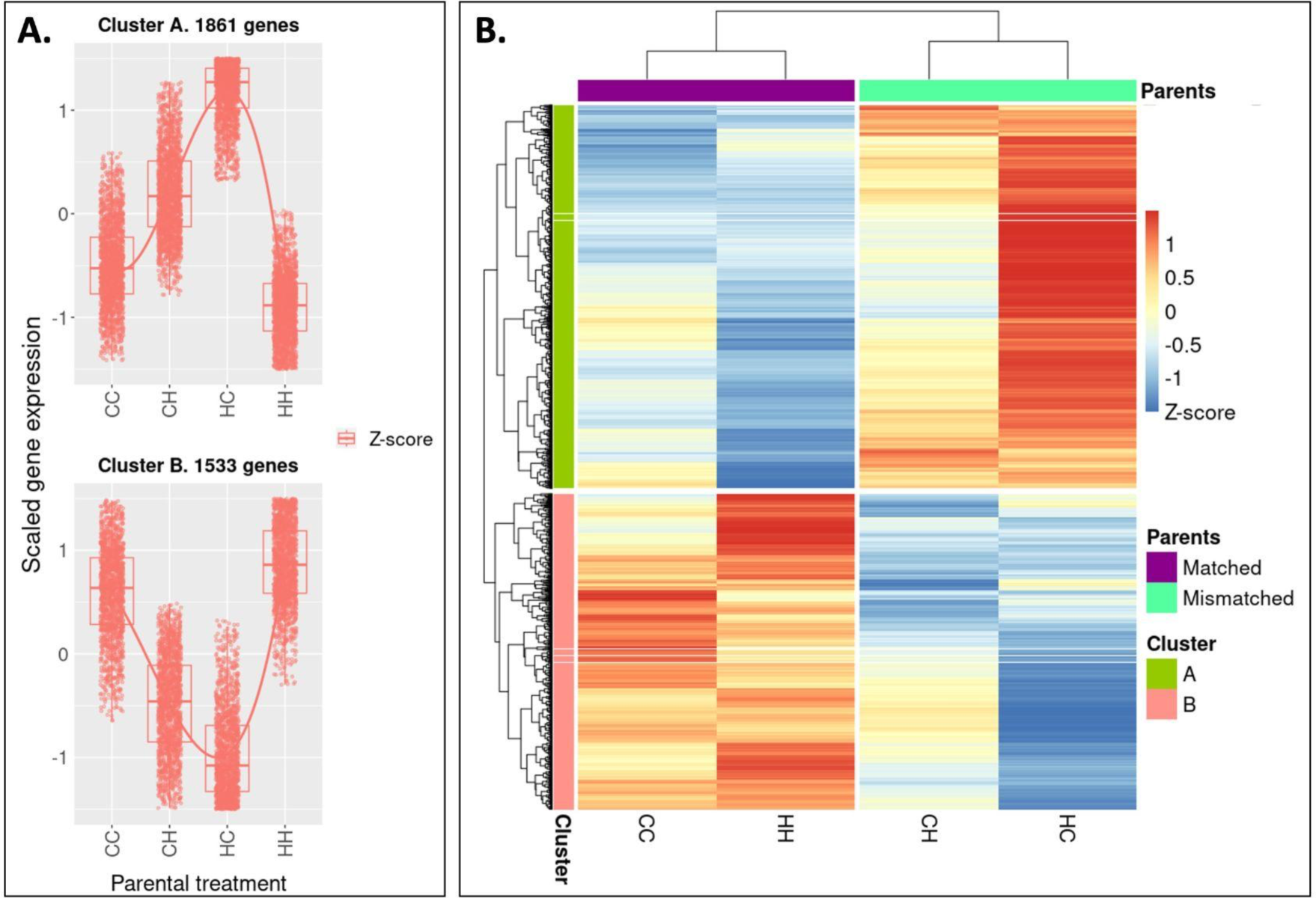
The two largest clusters of differentially expressed genes due to the interaction between parental treatments. DEGs were clustered based on their scaled expression profiles (Z-score). A) Box plots and median expression are shown for each parental treatment combination. B) Heatmap of Z-score values. In the parental treatment, the first letter stands for the paternal thermal environment, the second for the maternal one, where “C” = Control and “H” = +1.5°C.

Accordingly, pairwise comparisons between offspring from different parental thermal histories showed that HH offspring were more different to offspring from mismatched parents, rather than from CC offspring (Fig. 4). Overall, 601 HH genes were commonly DE and represent the molecular signature of biparental exposure to warming (Suppl. Fig. S1B; Suppl. Table S19). Among those, the upregulated genes are involved in ribosome biogenesis, ncRNA processing, especially rRNA and tRNA, gene expression and transcription. The downregulated ones, on the other hand, include nuclear steroid receptors, and several genes with oxidoreductase activity functions (Suppl. Table S20). Strikingly, there was little overlap between the three parental exposure combinations, with only 35 genes always differentially expressed in offspring with either one or both parents raised at elevated temperature compared to CC offspring (Suppl. Fig. S1A; Suppl. Table S21). Accordingly, distinct parental thermal histories differently shaped the offspring response to developmental warming, with mostly unique sets of genes expressed by offspring when raised at elevated temperature (Fig. 5). Interestingly, the smallest number of DEGs, 34, was found for the comparison between offspring raised at elevated temperature *versus* at control when born from HH parents (Suppl. Table S22). 26 of these genes were unique to this comparison, with functions related to oxidoreductase activity, iron ion homeostasis and heme biosynthesis.

## Discussion

Here we show that molecular responses to ocean warming in a coral reef fish depend on conserved and parent-specific effects. Surprisingly, paternal thermal exposure to elevated temperature had a stronger effect on differential gene expression in offspring compared with maternal exposure to elevated temperature. However, it was the consistency between the two parental experiences that was key to offspring acclimation potential. While different parental thermal histories uniquely shaped how offspring responded to warming during their own developmental period, these parental effects did not interact with the sex nor the environment of the offspring, and therefore had a carry-over rather than an anticipatory nature, contrasting with what has been found in some other fish species (Chang et al., 2021; Shama, Strobel, Mark, & Wegner, 2014). Strikingly, the smallest transcriptional response to warming experienced in the F2 generation was found in offspring from parents where both the mother and father were exposed to elevated temperature, suggesting a limited molecular response was enough for these offspring to acclimate to a warmer environment.

Among the differentially expressed genes due to the parental thermal experience, approximately half was exclusively due to either the father or the mother. Similarly, specific molecular signatures in *A. polyacanthus* brains were found to be maternally and paternally inherited in response to elevated CO_2_ exposure (Monroe, Schunter, Welch, Munday, & Ravasi, 2021). Parent-specific effects are usually expected when the reliability of the environmental information transmitted by one parent is greater than the other one (Bell & Hellmann, 2019; Burke, Nakagawa, & Bonduriansky, 2020). However, since *A. polyacanthus* is not sexually dimorphic, both sexes have the same ecology, and they provide joint parental care, these parent-specific effects are likely due to other differences between the two sexes, notably the way information is transmitted by sperm and oocyte (Bell & Hellmann, 2019; Perez & Lehner, 2019). For example, different epigenetic factors have been found to be independently transferred by mothers and fathers in sticklebacks following exposure to higher temperatures (Fellous, Wegner, John, Mark, & Shama, 2021). Moreover, despite parental effects have previously been primarily considered a female prerogative, the importance of paternal contributions has been increasingly recognized (Crean & Bonduriansky, 2014; Immler, 2018; Rando, 2012). In fact, we observed a larger effect of the paternal experience compared to the maternal one, as also seen in the marine tubeworm *Galeolaria caespitosa* exposed to warming (Guillaume, Monro, Marshall, & Pfrender, 2015). Therefore, our results show that 80-day old *A. polyacanthus* exhibit parent-specific intergenerational effects that influence distinct molecular pathways in offspring livers, and where the sperm-born information is the strongest driver of gene expression change.

The father-specific effects due to the paternal exposure to warming involved the activation of stress related pathways, like apoptosis and suppression of innate immunity response, as well as epigenetic gene expression regulatory mechanisms through histone modifications and microRNA (miRNA) mediated gene silencing. Chromatin modifications and miRNA production could represent epigenetic defense mechanisms against stress-induced genome mutations, potentially transferred from the male germline to the offspring (Immler, 2018). Differential histone regulation has been found to be influenced by the paternal phenotype in *A. polyacanthus* exposed to simulated ocean acidification (Monroe et al., 2021), and in stickleback sperm following warming (Fellous et al., 2021). Sperm-born miRNAs from stressed fathers, on the other hand, alters mice offspring behavior and metabolism (Gapp et al., 2014; Rodgers, Morgan, Bronson, Revello, & Bale, 2013), and stressed zebrafish show changes in sperm miRNA levels (Ord, Heath, Fazeli, & Watt, 2020). Here, when fathers alone were exposed to elevated temperature, their offspring additionally showed downregulation of several ribosomal proteins and translation related genes, indicating an inhibition of the protein synthesis machinery, likely because of cellular stress, and linked to trade-offs in energy investment (Sokolova, Frederich, Bagwe, Lannig, & Sukhotin, 2012; Spriggs, Bushell, & Willis, 2010). In support of these findings, these juveniles were shorter compared to offspring from other parental combinations (Spinks et al., 2022), suggesting impaired growth due to redirection of energy towards thermal stress response processes. Interestingly, offspring were also lighter compared to control, not only when the father alone was exposed to warming but also when both parents were. Notably, adipogenesis and fatty acid metabolism related genes were downregulated in offspring from both those parental combinations. The paternal exposure to warming therefore might interfere with offspring lipid storage and overall energy homeostasis, since lipid, and in particular fatty acids, are the main fish metabolic energetic source necessary for growth, reproduction and swimming (Tocher, 2003). Hence, the fathers’ development at elevated temperature causes the activation of pathways linked to stress in offspring, which result in the reallocation of energy expenditure from macromolecular biosynthesis and metabolism to repair mechanisms, with some pathways potentially epigenetically controlled.

Compared to the paternal effects, maternal thermal exposure appeared to be less influential to 80-day old *A. polyacanthus*, but likewise stressful, especially for offspring from pairs where only the mother experienced warming. Mothers’ development at elevated temperature mainly resulted in changes in offspring lipid and retinol metabolism, both linked to energy metabolism regulation (Klyuyeva et al., 2021). Apolipoproteins, for example, key regulators of cholesterol and triglyceride transport (Dominiczak & Caslake, 2011), were upregulated, especially when offspring from mother-only exposed pairs were raised at elevated temperature themselves. Transgenerational thermal acclimation through improved aerobic performance in *A. polyacanthus* has been linked to increases in lipid metabolism (Bernal et al., 2021; Veilleux et al., 2015), notably with the transgenerational upregulation of apolipoproteins (Veilleux et al., 2015). Our results therefore suggest that this metabolic adjustment, needed to sustain the increased energy demand at elevated temperatures, might primarily be a maternally derived effect. Despite these potentially beneficial effects, maternal exposure alone also caused downregulation of glutathione S-transferases, whose dysregulation is commonly associated with liver disease forms in mammals (Kirpich et al., 2011), and differential regulation of genes involved in immune and inflammatory responses, as well as hepatocyte apoptosis, indicating long-term detrimental effects caused by mothers’ exposure to warming. Therefore, similarly to the stress-related pathways elicited by paternal exposure, maternal experience of warming in *A. polyacanthus* causes some potentially maladaptive parental effects, especially when fathers were raised at control temperature. Nevertheless, it is possible that maternally influenced changes in lipid and energy metabolism and paternally inherited epigenetic stress-response mechanisms could represent adaptive parental effects, potentially responsible for the transgenerational acclimation to warming previously reported for this species (Bernal et al., 2021; Donelson et al., 2012). Metabolism adjustment might allow offspring to meet thermally induced rises in energy demand, while the activation of epigenetic repair mechanisms could help protect the integrity of the DNA at elevated temperatures. Hence, different traits appear to have independent acclimation potentials, and while transgenerational exposure to environmental stressors such as ocean warming might enhance tolerance to such stressor, this might come at the cost of reducing overall fitness, e.g., increasing disease susceptibility through immune response suppression or decreasing reproductive output. Ultimately, such trade-offs and the balance between costs and benefits of adaptive plasticity (DeWitt, Sih, & Wilson, 1998) will determine the organisms’ overall ability to adapt to environmental changes.

The exposure of both parents to elevated temperature resulted in the upregulation in their offspring of genes involved in gene expression regulation and production of regulatory RNAs responsible for protein synthesis. It also resulted in the smallest transcriptional response of offspring to their own experience of warming. Interestingly, increased peptide biosynthesis ability has been linked to transgenerationally acquired acclimation to warming in sticklebacks (Shama et al., 2016), while a reduced transcriptional response to thermal exposure could be a sign of increased tolerance to thermal stress. In this scenario, a small number of DEGs in offspring from parents that were both exposed to warming might indicate the lack of activation of pathways, e.g., genes related to stress response, not necessary to more acclimated individuals to respond to elevated temperature. Similarly, heightened tolerance to warming was associated with decreased gene expression plasticity across generations in the crustacean *Tigriopus californicus* (Kelly, Pankey, DeBiasse, & Plachetzki, 2017). Indeed, although a greater transcriptional response can sometimes imply higher plasticity and acclimation potential, studies have shown that increased tolerance to warming is often accompanied by reduced plasticity and gene expression changes in response to elevated temperature (Barley et al., 2021). Therefore, our results may indicate adaptive parental effects in offspring from parents that both experienced warming, as opposed to the negative parental effects found in offspring where one parent only was exposed to elevated temperature. Indeed, offspring from mismatched parents showed signs of endoplasmic reticulum stress and concomitant energy reallocation towards stress response and repair mechanisms. Moreover, downregulation of genes encoding for cytochrome C oxidase subunits suggest a reduced mitochondrial respiratory capacity for these offspring compared to controls and compared with offspring where both parents were exposed to elevated temperature. Swimming performance of offspring from this same experiment corroborates our molecular findings, with fish born from mismatched parents showing maladaptive swimming speed compared to controls and offspring from parents with a matched exposure to elevated temperature (Spinks, 2021). Such non-additive parental effects have also been seen in predator-induced sticklebacks, where offspring brain gene expression profiles differed depending if one or both parents had been predator-exposed (Hellmann et al., 2020), and single-parent effects on daughter mate choice were reversed when both parents were exposed to predation (Lehto & Tinghitella, 2020). Here, biparental exposure to warming is likely necessary for the transgenerational acclimation to increased temperature previously observed in this species (Bernal et al., 2021; Donelson et al., 2012; Veilleux et al., 2015). On the contrary, breeding pairs composed by individuals that developed at different thermal regimes, for example during a marine heatwave, will instead transfer to their offspring maladaptive stress condition signatures, which seems to override the beneficial adaptive benefits of parental exposure to warming.

Environmentally induced parental effects, if adaptive, might allow organisms to acclimate across generations to a rapidly warming world. Understanding the relative importance of fathers and mothers and how their thermal experiences are integrated in the offspring phenotype is crucial to make better predictions about species persistence potential. Here we show that warming induces a combination of shared, and, more strikingly, also father- and mother-specific molecular signatures, some of which likely beneficial to the offspring. Our results suggest that the key to transgenerational acclimation appears to lay in the consistency between the paternal and maternal thermal experiences. Exposure of both parents to elevated temperature caused increased expression of genes involved in transcriptional regulation and protein synthesis in the next generation, resulting in small adjustments needed in the offspring if raised at elevated temperature. However, if only one parent experienced warming, fish manifested energy trade-offs towards stress response and repair mechanisms, suggesting carry-over effects of a suboptimal condition of parents that developed at elevated temperature. Therefore, mismatched thermal experiences of mothers and fathers might interfere with the adaptive potential of parental effects. In the race between thermal acclimation and climate change, parental experience will likely play a crucial role. Ultimately, how beneficial parental effects are will depend on the balance between the costs and benefits of plasticity, and on the consistency between the past environmental experiences of mothers and fathers.

## Data accessibility

RNAseq data for all individuals can be found under the BioProject PRJNA998209.

## Authors’ contributions

L.C.B., J.M.D. and R.K.S. designed the experiment and collected the samples. L.C.B. and R.K.S. managed the fish rearing. L.C.B. prepared the samples for sequencing, analyzed the sequencing data and wrote the first draft of the manuscript, with input from C.S.. J.M.D., R.K.S., P.L.M. and T.R. secured the funding. All authors read, provided comments and gave final approval for publication.

## Acknowledgements

We would like to thank the Marine and Aquaculture Research Facilities Unit at JCU and KAUST Bioscience Core Lab for their help and support.

