## Supplementary Figures for "Matching maternal and paternal experiences underpin molecular thermal acclimation"

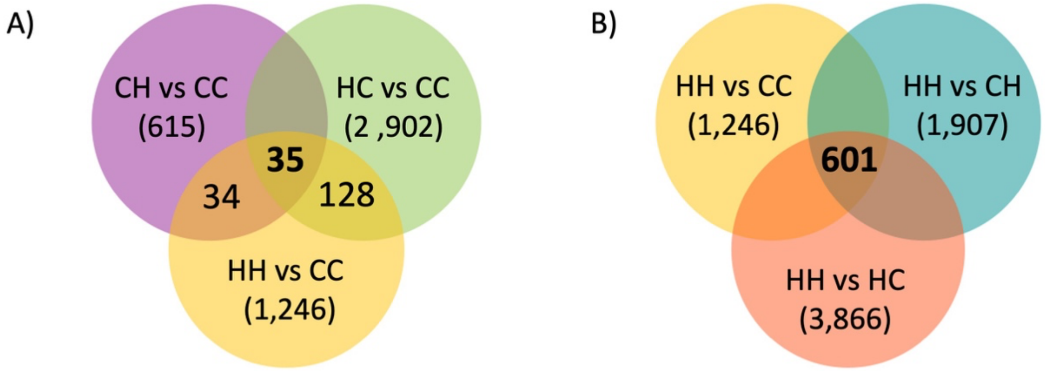


**Supplementary Figure S1.** Venn diagrams of differentially expressed genes in pairwise comparisons between A) CC; B) HH *versus* other parental thermal treatment offspring. The first letter stands for the paternal thermal environment, the second for the maternal one; "C" = control temperature, "H" = +1.5°C
